# Distinct glycosaminoglycan chain length and sulfation patterns required for cellular uptake of Tau, Aβ, and α-Synuclein

**DOI:** 10.1101/207035

**Authors:** Barbara E. Stopschinski, Brandon B. Holmes, Gregory M. Miller, Jaime Vaquer-Alicea, Linda C. Hsieh-Wilson, Marc I. Diamond

## Abstract

Transcellular propagation of aggregate “seeds” has been proposed to mediate progression of neurodegenerative diseases in tauopathies and α-synucleinopathies. We have previously determined that tau and α-synuclein aggregates bind heparan sulfate proteoglycans (HSPGs) on the cell surface. This mediates uptake and intracellular seeding. The specificity and mode of binding to HSPGs has been unknown. We used modified heparins to determine the size and sulfation requirements of glycosaminoglycan (GAGs) binding to aggregates in biochemical and cell uptake and seeding assays. Aggregates of tau require a precise GAG architecture with defined sulfate moieties in the N- and 6-O-positions, whereas α-synuclein and Aβ rely slightly more on overall charge on the GAGs. To determine the genetic requirements for aggregate uptake, we individually knocked out the major genes of the HSPG synthesis pathway using CRISPR/Cas9 in HEK293T cells. Knockout of EXT1, EXT2 and EXTL3, N-sulfotransferase (NDST1), and 6-O-sulfotransferase (HS6ST2) significantly reduced tau uptake. α-Synuclein was not sensitive to HS6ST2 knockout. Good correlation between pharmacologic and genetic manipulation of GAG binding by tau and α-synuclein indicates specificity that may help elucidate a path to mechanism-based inhibition of transcellular propagation of pathology.

## Introduction

Tau and α-synuclein aggregates bind heparan sulfate proteoglycans (HSPGs) on the cell surface to mediate uptake and intracellular seeding (1). However, the specificity of this interaction is unknown. Heparan sulfate (HS) is a linear glycosaminoglycan (GAG) composed of disaccharide repeats with sulfate moieties at discrete N-, C2-, C3-, and C6-residues (Figure 1), and is synthesized in all known eukaryotic cells (2,3). During synthesis, HS chains are covalently attached to core proteins, forming heparan sulfate proteoglycans (HSPGs). Many HSPGs are secreted into the extracellular matrix or membrane associated, either via a transmembrane domain or a glycosyl-phosphatidylinositol (GPI) anchor (2,3). The structure of the HS chains is determined by the activity of synthetic enzymes, availability of precursor molecules, and flux through the Golgi apparatus. The structural diversity of HSPGs among different cell and tissue types accounts for their interaction with myriad proteins. The so-called *heparan sulfate interactome* includes proteins involved in cell attachment, migration, invasion and differentiation, morphogenesis, organogenesis, blood coagulation, lipid metabolism, inflammation, and injury response (2,4). Some HS-binding proteins bind HS carbohydrates with great specificity. The interaction between HSPGs and their binding proteins can depend on modification, domain, charge and sugar conformation (2,5,6). Some interactions are based on defined chemistry. For example the binding of a protein may require spatially separated patches of sulfated residues on the GAGs. Purely charged-based interactions are more flexible, as they are based on electrostatic interactions between the positively charged residues in the binding protein and the negatively charged sulfate residues in the HS chain (2). We do not know whether the interactions of tau and α-synuclein with HSPGs are specific to sulfation positioning, or are purely based on charge density.

**Figure 1:**
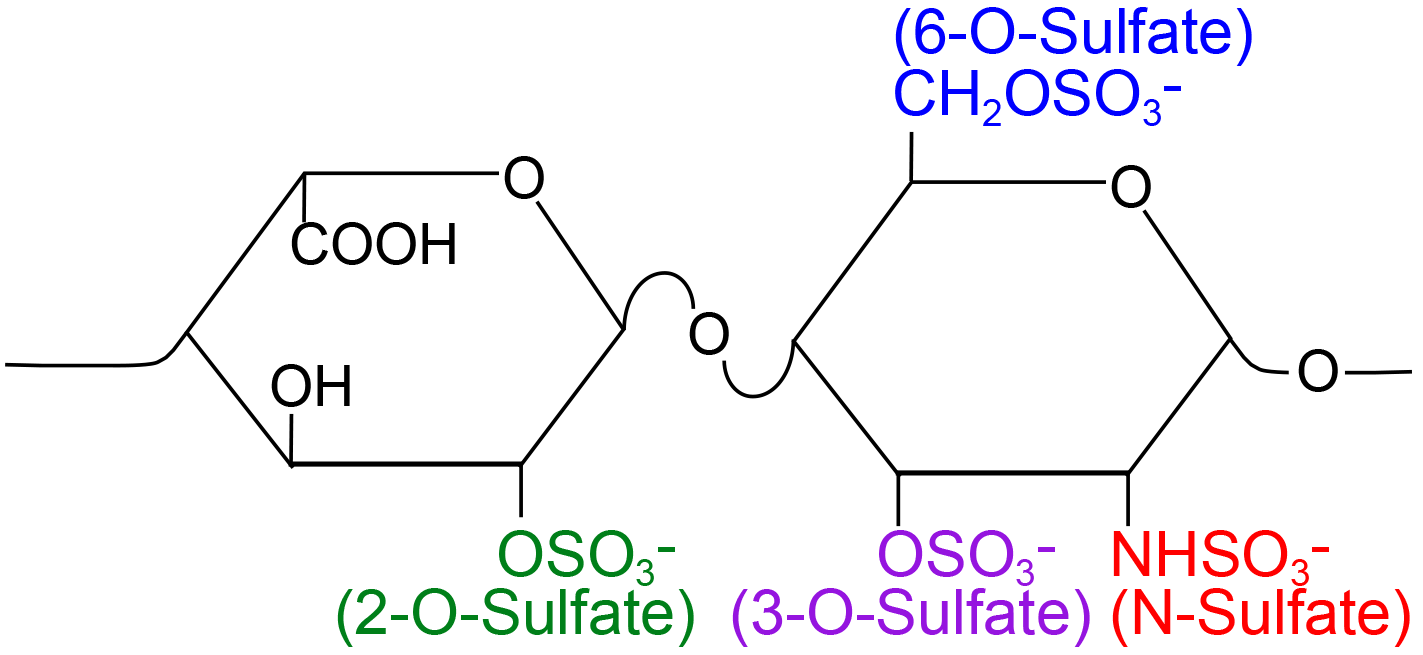
The basic disaccharide unit of heparin. Uronic acids (left) are sulfated at the 2-O position while glucosamine residues (right) are sulfated at the N-, 3-O-, and 6-O-positions.

We previously observed that exogenously applied tau and α-synuclein aggregates bind to HSPGs on the cell surface to trigger cellular uptake and induce intracellular seeding. Thus, the HSPG:aggregate interaction may represent a critical step for the spread of pathology in neurodegenerative diseases (1). Other laboratories have similarly observed that Aβ and prion protein bind to HSPGs on the cell surface to trigger their internalization (7–11). It is not clear which HSPG properties drive the interaction with aggregates, and if the required GAG composition differs among amyloid proteins.

We analyzed the GAG binding requirements for recombinant tau, α-synuclein, and Aβ amyloid fibrils. We used a small heparin mimetic library to determine the critical size and sulfation requirements for HS to bind to aggregates in biochemical and cell-based assays, and used CRISPR/Cas9 knockout to test specific components of the HSPG synthesis pathway. Binding of tau fibrils to HSPGs requires modification-specific interactions, whereas α-synuclein and Aβ fibrils rely more on overall sulfation rather than specific sulfate moieties.

## Results

### Carbohydrate microarrays identify unique binding patterns for amyloid proteins

Carbohydrate microarrays allow rapid analysis of interactions between proteins and GAGs (12). We used this approach to investigate the structural features of heparin that are required for binding to tau, α-synuclein, Aβ and huntingtin seeds (Figure 2). As for prior studies (13–15), we used a library of modified heparin polysaccharides (Neoparin, Alameda, CA) to investigate sulfation requirements for binding: heparin (Hep), N-desulfated heparin (De-N), 6-O-desulfated (De-6-O), 2-O-desulfated (De-2-O), O-desulfated heparin (De-O), fully desulfated heparin (De-S, in which >90% of all sulfates are removed) and over sulfated heparin (Over-S, in which there are >3.5 sulfates per disaccharide unit).

**Figure 2:**
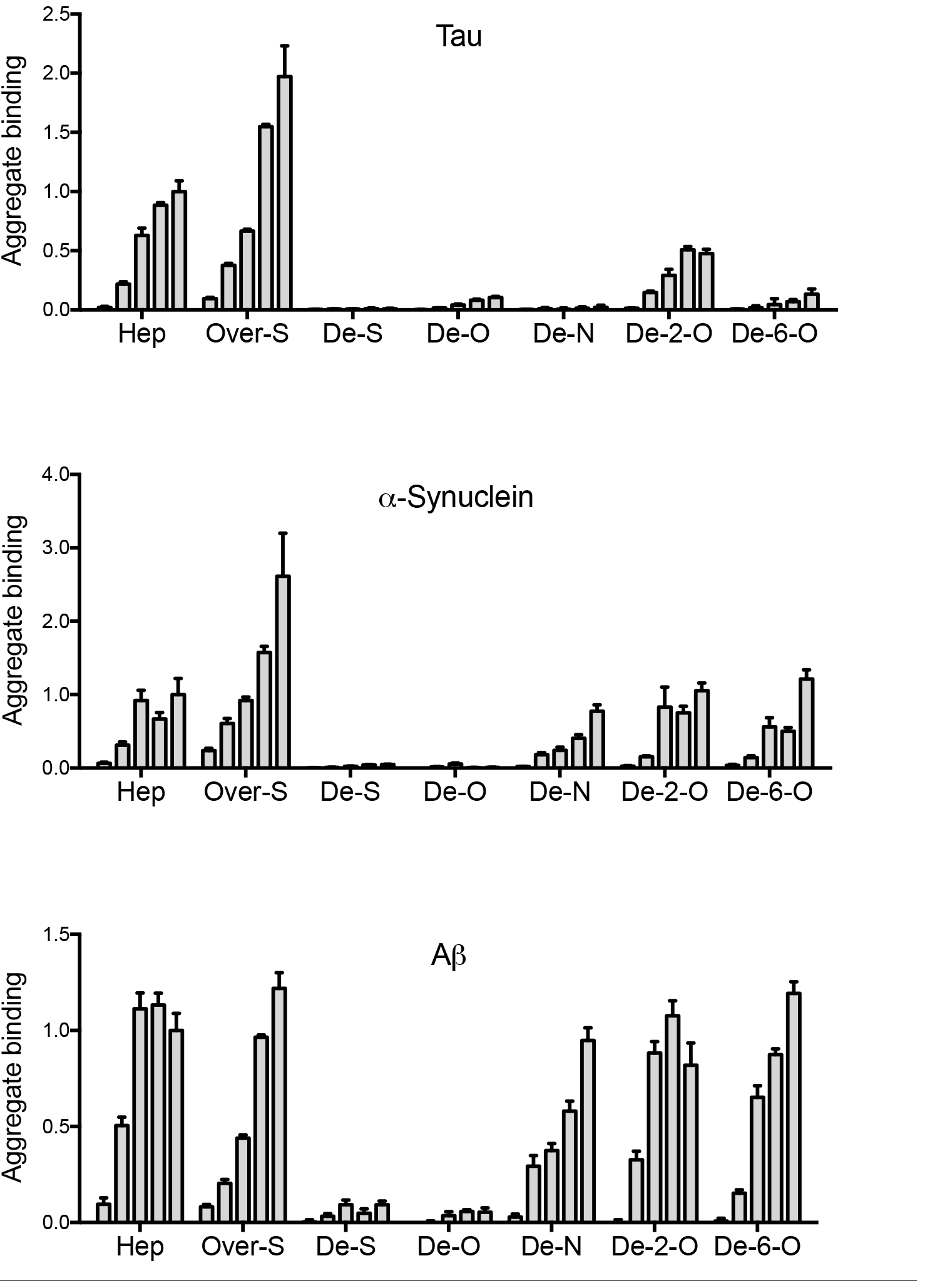
Carbohydrate microarray analyses identify unique amyloid/GAG interactions. Biotin-labeled tau, α-synuclein, Aβ fibrils at 1μM (monomer equivalent) were applied to a glass surface coated with heparin derivatives at 0.5, 1, 5, 10, and 15 μM (left to right): heparin (Hep), over-sulfated heparin (Over-S), fully desulfated heparin (De-S), O-desulfated heparin (De-O), N-desulfated heparin (De-N), 6-O-desulfated (De-6-O), 2-O-desulfated (De-2-O). We measured relative fibril binding with fluorescently tagged streptavidin, averaging 10 spots in each given carbohydrate concentration. Note distinct binding patterns for De-N, De-2-O, and De-6-O heparins. Each sample was analyzed in triplicate. Data represent an average value for 10 spots at a given carbohydrate concentration. Error bars show SEM.

We applied nanoliter volumes of the heparins at concentrations from 0.5 μM to 15 μM on glass microarray surfaces coated with poly-L-lysine. We then applied biotinylated full-length tau, α-synuclein, Aβ42, and huntingtin exon 1 (HttExon1Q50) fibrils to the microarray, and visualized the bound proteins with an anti-biotin antibody tagged with Cy5. Tau, α-synuclein, and Aβ aggregates bound heparin in a concentration-dependent manner (Figure 2). Huntingtin fibrils exhibited no binding (data not shown) and were not analyzed further. Desulfated heparin did not bind any of the fibrils, suggesting that sulfation is a critical component of the aggregate-GAG interaction (Figure 2). Our results comported with previous reports that tau, α-synuclein, and Aβ, but not Htt, are heparin-binding proteins (1,7,16,17).

The different seeds exhibited unique sulfation requirements for binding. Tau efficiently bound heparin and 2-O-desulfated heparin, whereas 6-O-desulfation, N-desulfation, or O-desulfation abolished its binding (Figure 2). α-Synuclein and Aβ fibrils each required O-sulfation for heparin binding. However, the removal of N-, 6-O, or 2-O sulfation did not significantly inhibit their binding (Figure 2). Thus, for α-synuclein and Aβ fibrils, no single sulfate moiety was required to mediate the GAG interaction.

### Inhibition of amyloid uptake requires specific heparin sulfation pattern

Our group and others previously observed that heparin inhibits the cellular binding and uptake of tau, α-synuclein, and Aβ (1, 7). Heparin directly interacts with the HS binding domains and prevents seed binding to HSPGs on the cell surface. Thus we tested whether the structural determinants of the seed-GAG interaction observed in the microarray assay would translate to seed internalization and propagation in cells.

We labeled tau, α-synuclein, and Aβ fibrils with a fluorescent Alexa Fluor 647 dye and applied them to C17.2 cells in culture for 4 hours (tau and α-synuclein) or 20 hours (Aβ). We measured aggregate uptake with flow cytometry by quantifying the median fluorescence intensity per cell (MFI). Heparin decreased seed internalization dose-dependently for all 3 fibril types (Figure 3).

**Figure 3:**
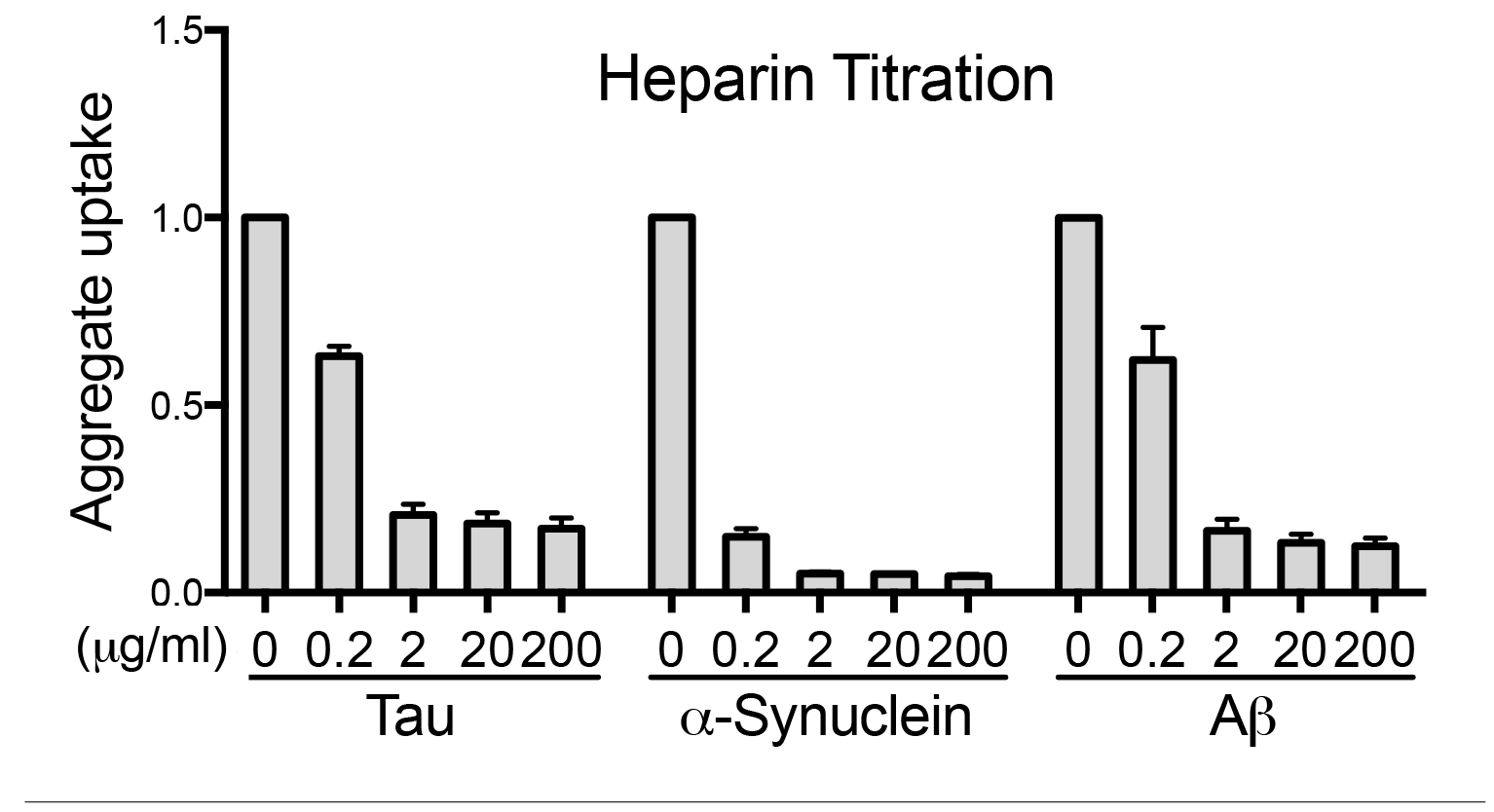
Heparin blocks aggregate uptake of Tau, Synuclein and Aβ. Tau, α-synuclein, or Aβ seeds labeled with Alexa Fluor 647 fluorescent dye were applied to cells with increasing doses of heparin (0.2, 2, 20 and 200 μg/ml). We quantified aggregate uptake based on the median fluorescence intensity (MFI) per cell by flow cytometry. Heparin dose-dependently decreased cellular uptake in all cases. Each condition was recorded in triplicate, and values represent the average of three separate experiments. Data reflects uptake relative to the untreated samples. Error bars = SD.

We next tested the desulfated heparins as inhibitors of aggregate internalization (Figure 4). Tau aggregate uptake was strongly inhibited by 2-O-desulfated heparin, similar to standard heparin. Thus, removal of 2-O-sulfates did not disrupt tau uptake inhibition. The inhibitory potency of 6-O desulfated heparin was reduced compared to 2-O-desulfated heparin and standard heparin, whereas N-desulfated heparin had virtually no effect (Figure 4). In summary, as for heparin binding *in vitro*, competitors for tau uptake in cells require 6-O and N-sulfation, while 2-O-sulfation is dispensable.

**Figure 4:**
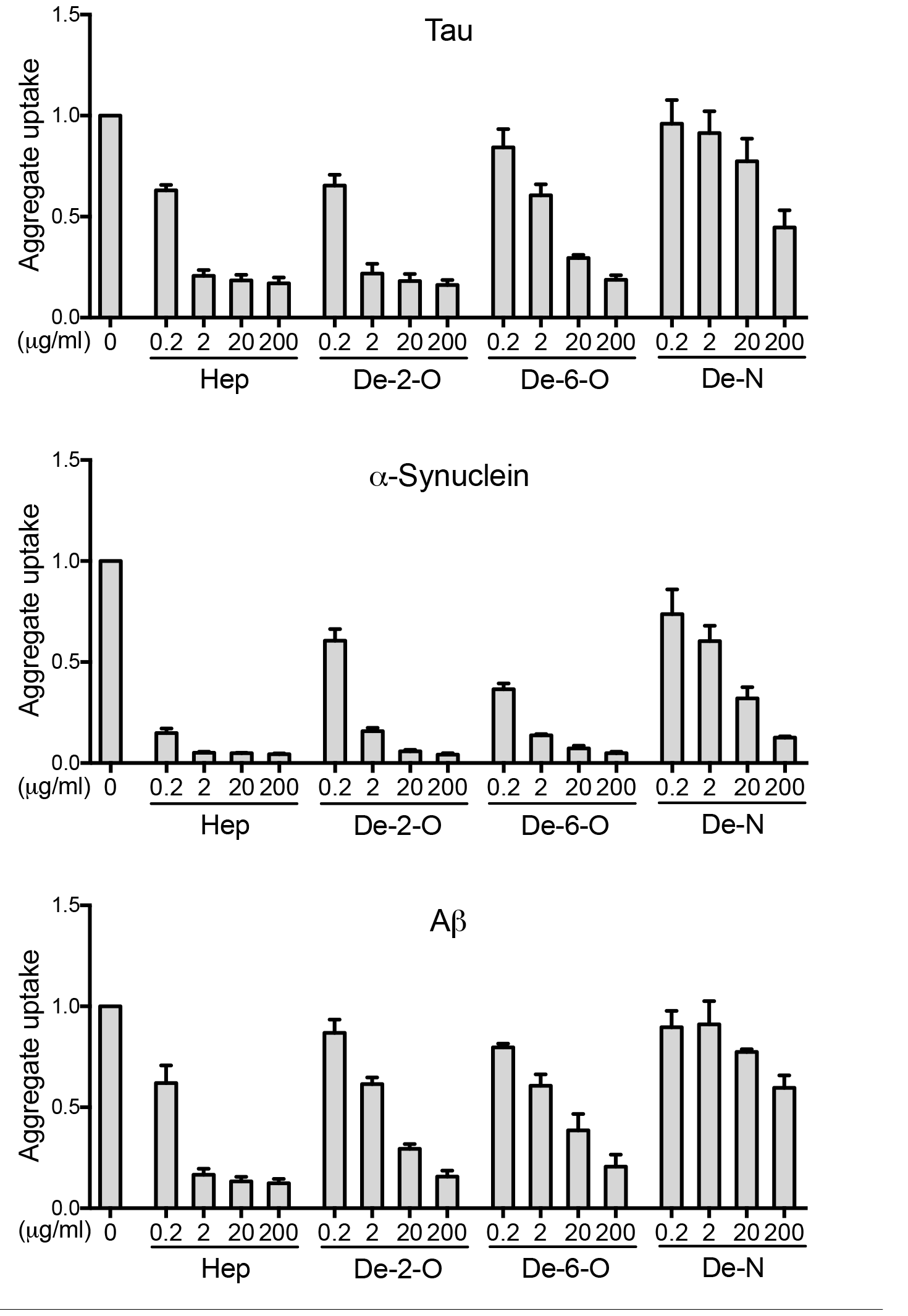
Heparin-mediated inhibition of aggregate uptake requires specific sulfation patterns. Normal heparin and 2-O, 6-O, and N-desulfated forms were tested as inhibitors of cellular uptake for tau, α-synuclein, and Aβ aggregates. N- and 6-O-desulfated heparins weakly inhibited, whereas 2-O-desulfated heparins strongly inhibited tau uptake. For α-synuclein and Aβ, removal of any of the 3 sulfate moieties reduced inhibitor efficacy. Tau uptake thus depends strongly on 6-O and N-sulfation to bind HSPGs. Each condition was recorded in triplicate, and values represent the average of three separate experiments. Data reflect uptake relative to the untreated group. Error bars show SD.

The structural requirements differed for the inhibition of α-synuclein and Aβ (Figure 4). Compared to standard heparin, removal of N-sulfation strongly reduced the inhibitory activity of the compounds on aggregate uptake. However, the removal of 6-O or 2-O sulfation from heparin also reduced its inhibitory effect on α-synuclein and Aβ uptake in comparison to unmodified heparin. Thus, N-sulfation may be critical, but other sulfates also contribute to aggregate binding. Compared to tau, inhibition of α-synuclein and Aβ internalization requires overall sulfation as opposed to sulfate moieties at specific residues.

### Inhibition of aggregate uptake requires a critical chain length

To determine the role of polysaccharide chain length, we tested the inhibitory activity of fractionated heparin moieties of various chain lengths in the cell uptake assay. Heparin fragments composed of 4, 8, 12, and 16 sugar molecules (referred to as 4-mer, 8-mer, 12-mer and 16-mer respectively) were individually incubated with labeled aggregates, applied to C17.2 cells, and tested for inhibition of uptake (Figure 5). For tau, shorter heparin fragments (4-mer and 8-mer) had no activity, while the longer fragments (12-mer and 16-mer) modestly inhibited uptake (Figure 5). No compound had a potency similar to heparin (~50-mer), even when taking into account the number of individual saccharide units. We conclude that larger heparin chains are necessary to inhibit tau uptake.

**Figure 5:**
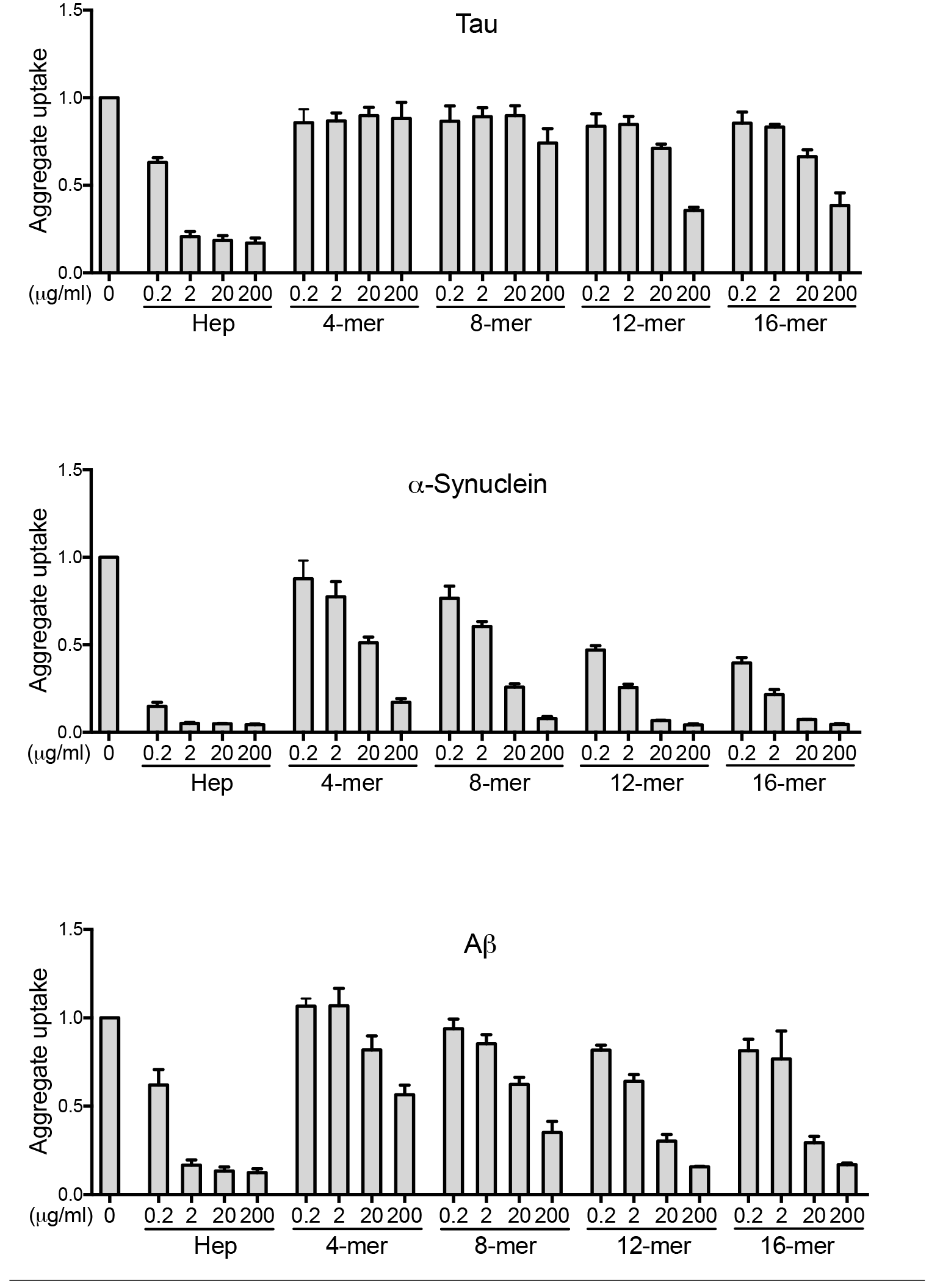
Inhibition of aggregate uptake requires a critical heparin size. 4-mer, 8-mer, 12-mer, and 16-mer heparins were evaluated as inhibitors of tau, α-synuclein, and Aβ internalization. Potency increased with the GAG chain length. Each condition was recorded in triplicate, and values represent the average of three separate experiments. Data reflect uptake relative to the untreated group. Error bars show SD.

Aβ fibrils exhibited greater sensitivity to shorter polysaccharides, and 12-mer and 16-mer inhibited uptake. As for tau, the uptake inhibition of Aβ increased with the chain length of the heparin. α-Synuclein aggregates, however, were dose-dependently inhibited by all fractionated heparins (Figure 5). Thus, depending on their target, heparins must consist of critical and distinct chain lengths to function as uptake inhibitors. We conclude that specific structural determinants of heparins, including sulfation pattern and size, govern their inhibitory activities for tau, α-synuclein, and Aβ aggregate uptake.

### Structural requirements for inhibition of seeding

Amyloid aggregates could gain entry to cells by multiple mechanisms, some of which could lead to seeding, and others not. Thus we tested heparins in an established seeding assay that consists of a monoclonal “biosensor” cell line that stably expresses tau repeat domain (RD) harboring the disease-associated mutation P301S, fused to yellow or cyan fluorescent proteins (RD-CFP/YFP) (18, 19). Upon binding to the cell surface, tau seeds trigger their own internalization, and induce intracellular aggregation of RD-CFP/YFP, enabling fluorescence resonance energy transfer (FRET). We use flow cytometry to quantify the number of cells exhibiting FRET. An α-synuclein biosensor that expresses full-length α-synuclein with the disease-associated mutation A53T tagged to either CFP or YFP (syn-CFP/YFP) functions similarly (19). Of note, we did not test seeding for Aβ due to the lack of a functional biosensor cell line. We incubated tau or α-synuclein seeds with heparins overnight, prior to direct exposure of the biosensor cells and incubation for 48 hours, followed by flow cytometry. To improve yield (due to low seeding efficiency) we re-exposed the α-synuclein biosensor cell line to seed/heparin complexes after passaging for an additional 48 hours prior to flow cytometry. Simultaneous application of heparin with tau and α-synuclein fibrils to the biosensor cell lines reduced seeding dose-dependently (Figure 6).

**Figure 6:**
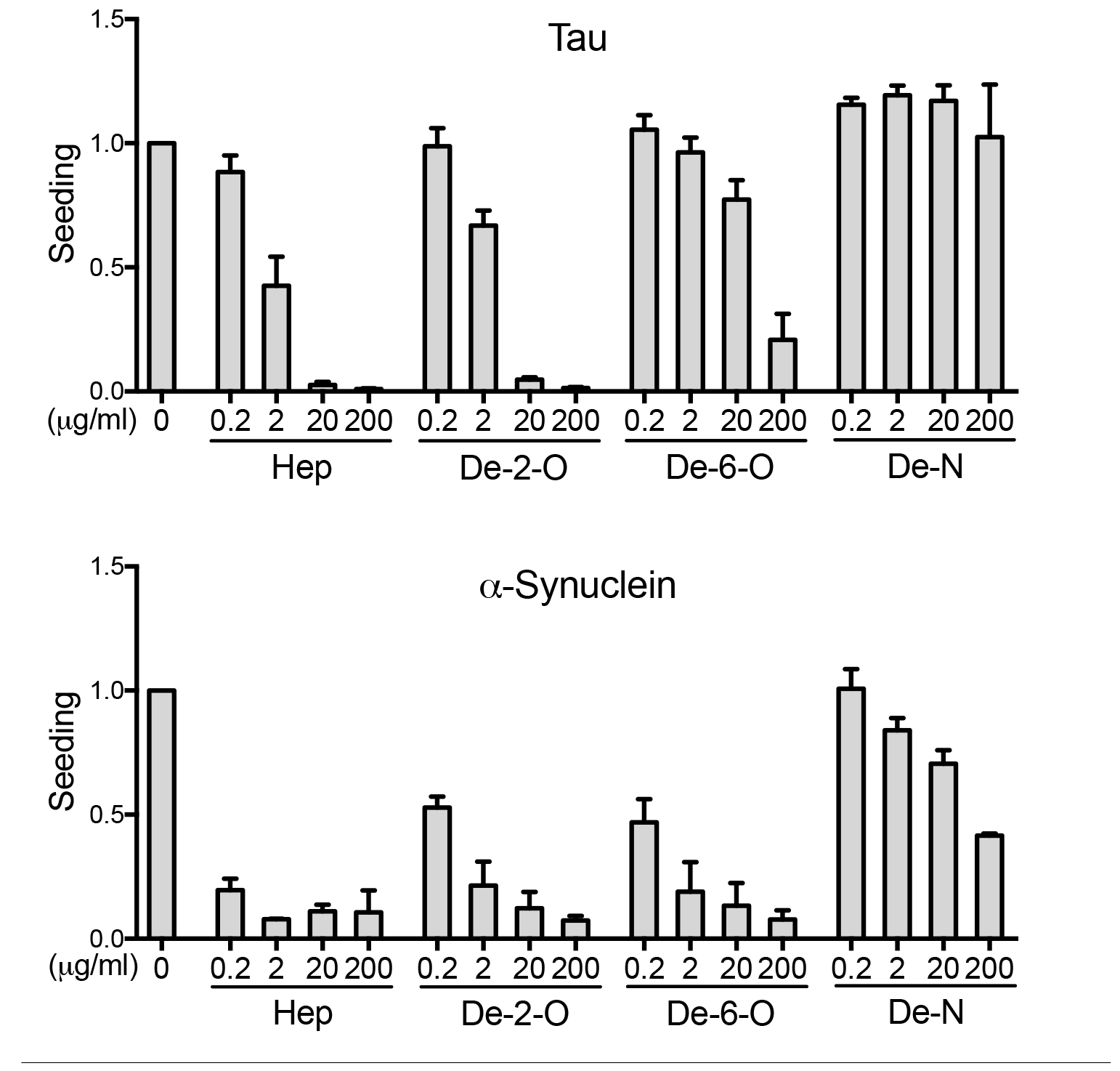
Sulfation pattern specifies inhibition of seeding. 2-O, 6-O, and N-desulfated heparins were evaluated as inhibitors against tau and α-synuclein seeding in biosensor cell lines. We used flow cytometry to measure seeding by quantifying the percentage of FRET-positive cells. Inhibition of tau seeding requires N-sulfation and, to a lesser extent, 6-Osulfation. By contrast α-synuclein primarily requires N-sulfation. In every experiment, each condition was tested in triplicate. Values represent the average of three separate experiments for tau, and two separate experiments for α-synuclein. The data reflect seeding relative to the untreated group. Error bars show SD.

We next applied the desulfated heparins to the seeding assay (Figure 6). 2-O-desulfated heparin blocked tau seeding almost as well as heparin, whereas 6-O-desulfated heparin lost most and N-desulfated heparin lost all activity against tau seeding. These observations were consistent with our prior finding that N-sulfation and 6-O-sulfation are important for cellular tau binding and internalization, whereas 2-O-sulfation is not. For α-synuclein, N-desulfated heparin lost most of its activity, concordant with results from our uptake studies. 6-O-desulfated heparin and 2-O-desulfated heparin also had reduced inhibitory activity compared to standard heparin. This indicated that overall sulfation rather than 6-O or 2-O-sulfation in particular is required to inhibit α-synuclein seeding.

We next tested fractionated heparins (4-mer, 8-mer, 12-mer, 16-mer). We observed no appreciable inhibitory activity against tau and only slight inhibitory activity against α-synuclein with shorter heparin fragments (4-mer and 8-mer), and weak activity with longer fragments (12-mer and 16-mer) for both fibril types. The results were qualitatively similar to the uptake assay, indicating that the inhibitory activity of the heparins against seeding increases with the saccharide chain length (Figure 7).

**Figure 7:**
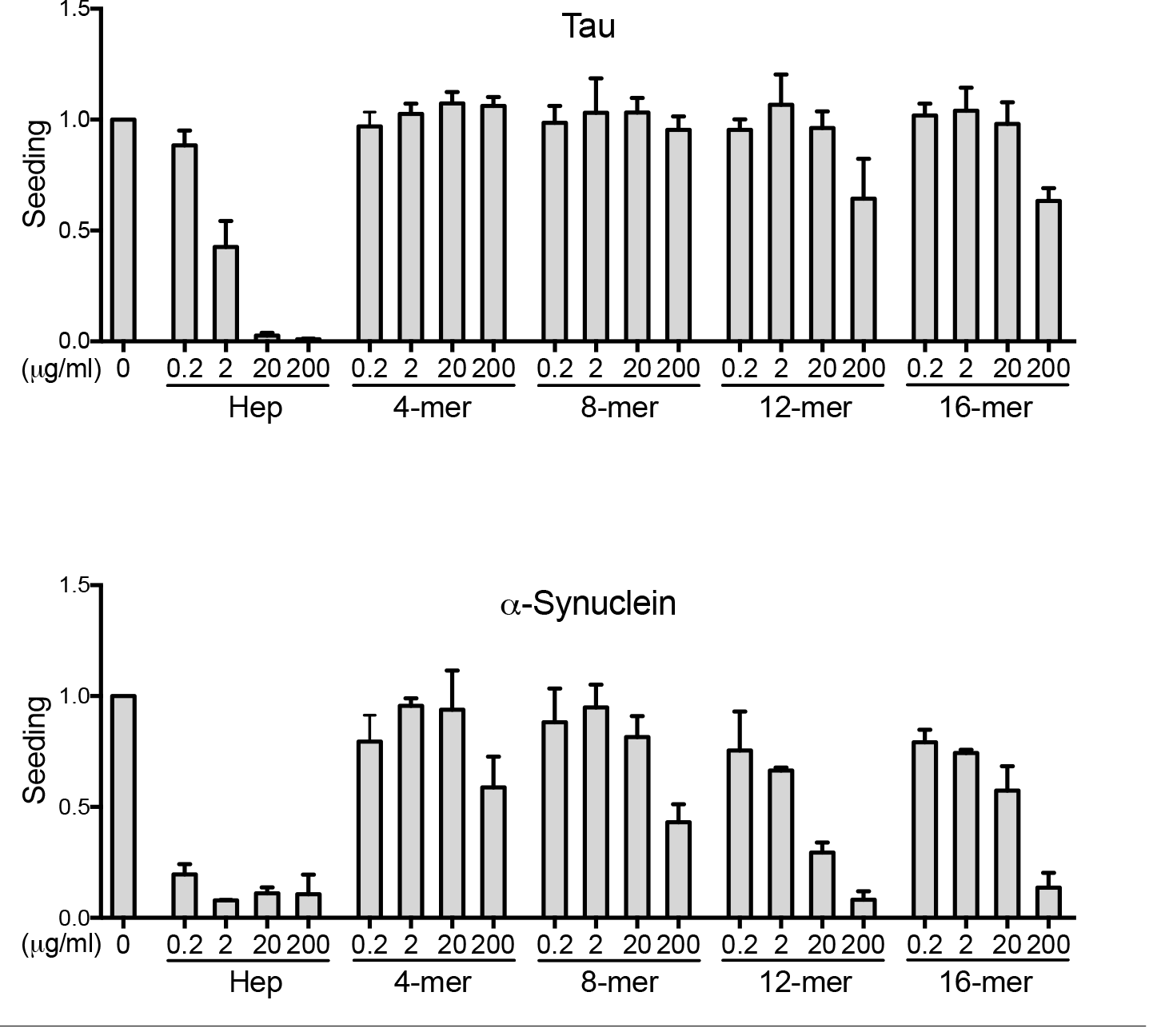
GAG size specifies inhibition of seeding. The inhibitory potency of different heparins against seeding of α-synuclein and tau depends on the chain lengths of the heparins. 4-mer, 8-mer, 12-mer, and 16-mer heparins were evaluated as inhibitors of tau and α-synuclein seeding. The inhibitory potency of the heparin fragments increased with GAG chain length. In every experiment, each condition was tested in triplicate. Values represent the average of three separate experiments for tau, and two separate experiments for α-synuclein. Data reflect seeding relative to the untreated group. Error bars show SD.

### HSPG synthetic genes required for uptake of aggregates

The HSPG synthesis pathway is a complex hierarchical cascade involving approximately 30 enzymes. After initial elongation, the carbohydrate chain undergoes a series of maturation steps in the Golgi apparatus. First, N-acetylglucosamine residues are N-deacetylated, followed by N-sulfation. Next, GlcA residues are epimerized to iduronic acid (IdoA). Finally, sulfate groups are attached to C3 and C6 of the GlcNAc residues and C2 of IdoA (2,3). We thus tested specific genes in the pathway to elucidate the structural determinants of cell surface HSPGs, and to further test the requirement of specific sulfation patterns for aggregate internalization.

We used CRISPR/Cas9 to individually knock out all the genes in the HSPG synthesis pathway and screen for inhibition of seed internalization in HEK293T cells (Table 1). We selected 5-6 separate human gRNAs for each gene and cloned them into the lentiCRISPRv2 vector (20). Plasmids containing the gRNAs for each gene were pooled and used for lentivirus production. Cells were exposed to lentivirus cultured for at least 10 days under puromycin selection prior to phenotypic screening. As a positive control, we included the gRNAs to target the valosin-containing peptide (VCP) gene, since the knockout of this gene is lethal in mammalian cells. In addition, we transduced the available gRNAs individually to identify the functional gRNAs for selected genes. We then used a single gRNA to produce polyclonal knockout cell lines and confirmed the presence of indels in the predicted DNA regions by Tracking of Indels by Decomposition (TIDE) (21).

**Table 1:** Genes included in the genetic screen. Major genes involved in the synthesis of HSPGs were screened for effects on tau and α-synuclein uptake. We also tested genes coding for proteoglycan core proteins (glypicans, syndecans) and SULF1/2 which diminish HSPG sulfation via arylsulfatase and endoglucosamine-6-sulfatase activity. Genes involved in the synthesis of the chondroitin proteoglycan (CSPG) synthesis pathway, as well as SQSTM1 were targeted as negative controls. To confirm gene knockout we targeted VCP, whose knockout is lethal after 5 days.

| Elongation | Modification | CSPG | Others |
| --- | --- | --- | --- |
| EXT1 | NDST1 | GALNACT1 | SULF1 |
| EXT2 | NDST2 | GALNACT2 | SULF2 |
| EXTL1 | NDST3 | CHSY | GPC2 |
| EXTL2 | NDST4 |  | GPC4 |
| EXTL3 | HS2ST1 |  | GPC6 |
|  | HS3ST1 |  | SDC1-4 |
|  | HS3ST2 |  |  |
|  | HS3ST3A1 |  |  |
|  | HS3ST3B1 |  | Controls |
|  | HS3ST4 |  | Scrambled |
|  | HS3ST5 |  | SQSTM1 |
|  | HS3ST6 |  | VCP |
|  | HS6ST1 |  |  |
|  | HS6ST2 |  |  |
|  | HS6ST3 |  |  |
|  | HPSE |  |  |

The knockout of five genes strongly inhibited tau uptake: EXT1, EXT2, EXTL3, NDST1 and HS6ST2. Four of these five genes (EXT1, EXT2, EXTL3, NDST1) also inhibited α-synuclein uptake (Figure 8, Table 2, Suppl. Figure 1). We had previously observed that EXT1 is required for cellular uptake of tau and α-synuclein aggregates (1). EXT1 is a glycosyltransferase that polymerizes heparan sulfate chains, and knockout of this gene reduces HSPG expression but does not affect other proteoglycan subtypes (i.e., chondroitin and dermatan sulfate proteoglycans) (22). EXT1 and EXT2 are co-polymerases, and both are required for proper HS chain elongation *in vivo* (23). EXTL3 is likewise a glycosyltransferase involved in the initiation and the elongation of the HS chain, and reduced levels create longer HS with fewer side chains (23).

**Figure 8:**
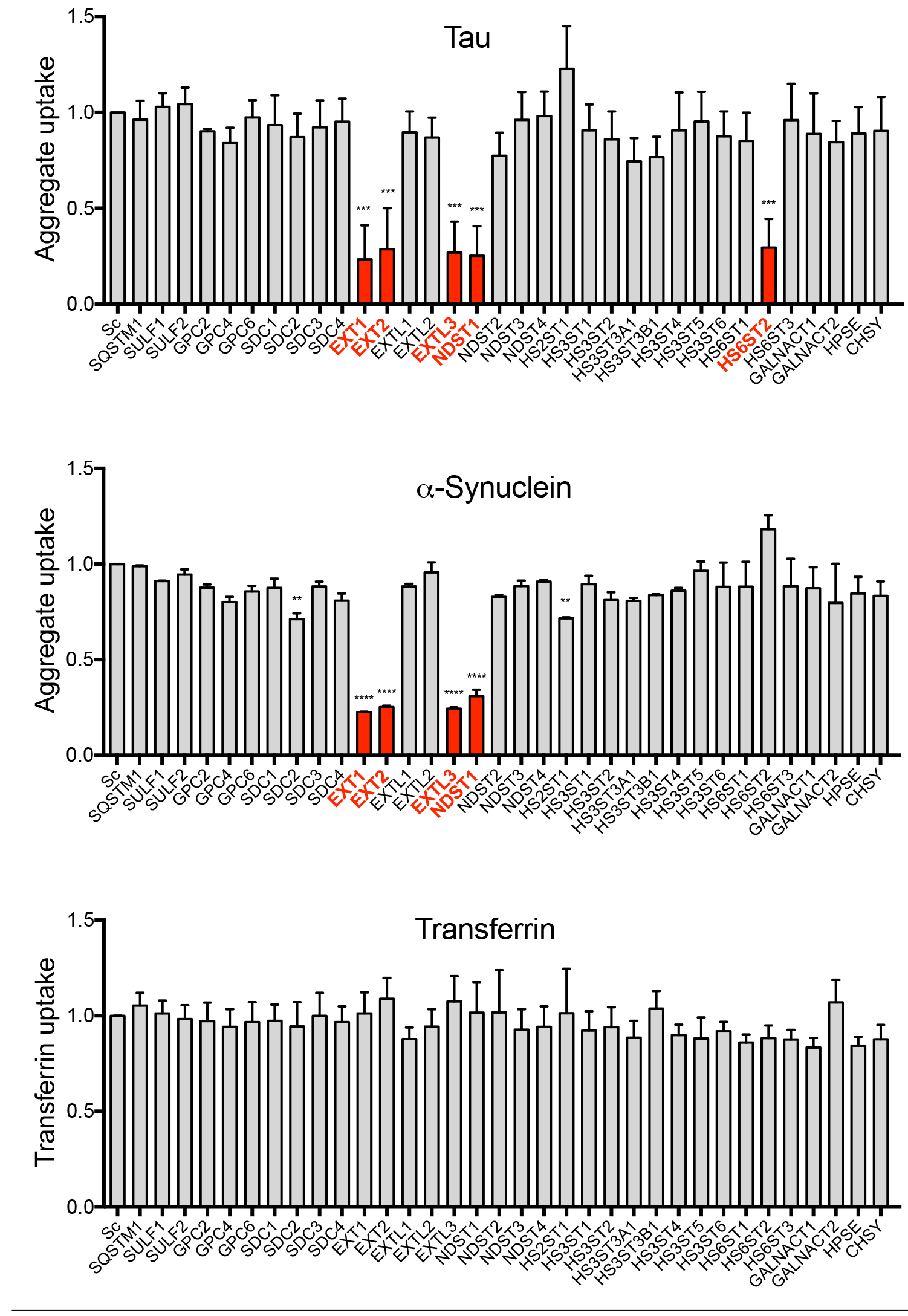
HSPG genes critical for the internalization of tau and α-synuclein aggregates. Genes implicated in HSPG synthesis were individually targeted in HEK293T cells using CRISPR/Cas9 to create polyclonal knockout lines. The cell lines were then tested for internalization of fluorescently labeled tau and α-synuclein aggregates by measuring median fluorescence intensity (MFI) per cell with flow cytometry. The cells were treated in parallel with fluorescently labeled transferrin to control for reduction of clathrin-mediated uptake. Knockout of four genes (Ext1, Ext2, EXTL3 and NDST1) reduced both tau and α-synuclein uptake, whereas the knockout of HS6ST2 only reduced tau uptake. None of the gene knockouts reduced transferrin uptake. Data was collected on 2 different days in duplicates for each cell line and each aggregate type, and normalized to uptake from control cells treated with scrambled gRNA (Sc). For transferrin uptake, the data from all 4 experimental days was combined. Red columns indicate the gene knockouts with the strongest effects. Error bars show SD. For statistical analyses, we combined the averages for each experimental day. ****P < 0.0001, ***P < 0.001, **P < 0.01; One-way Anova, Dunnett.

**Table 2:** Genes required for tau and α-synuclein uptake. The knockout of the listed genes reduced aggregate uptake by >50% and was statistically significant with p < 0.001; One-way Anova, Dunnett.

| TAU | α-SYNUCLEIN | Gene function |
| --- | --- | --- |
| Ext1 | Ext1 | Chain initiation/ elongation |
| Ext2 | Ext2 | Chain initiation/ elongation |
| EXTL3 | EXTL3 | Chain initiation/ elongation |
| NDST1 | NDST1 | N-sulfation |
| HS6ST2 | ---- | 6-O-sulfation |

NDST1 and HS6ST2 knockout also substantially reduced tau aggregate internalization (Figure 8, Suppl. Figure 1A). NDST1 places N-sulfate groups on glucosamine residues, while HS6ST2 attaches sulfate groups to 6-O-sulfoglucosamine residues. Thus, N- and 6-O sulfate moieties appear critical for HSPG-mediated uptake of tau aggregates. On the contrary, knockout of genes coding for 2-O-and 3-O-sulfotransferases had no effect (e.g. HS2ST1, HS3ST3A1) on tau uptake. These data indicate that N- and 6-O-sulfation are essential for tau uptake, corroborating our *in vitro* studies. Importantly, gRNAs encoding CSPG synthesis enzymes (CSGALNACT and CHSY) did not affect tau uptake (Figure 8). For α-synuclein, only knockout of NDST1 sulfotransferase strongly reduced aggregate uptake (Figure 8 and Suppl. Figure 1B). Knockout of other sulfotransferases (e.g. HS2ST1) mildly reduced uptake or had no effect (Figure 8). Consistent with the *in vitro* studies, enzymes responsible for N-sulfation were important for α-synuclein uptake, whereas those responsible for sulfation at other positions did not play a critical role. No gene knockouts reduced the internalization of labeled transferrin, which is controlled by clathrin-mediated endocytosis, suggesting lack of non-specific inhibitory activity on endocytosis (Figure 8).

To confirm specificity of our knockouts, we rescued the main genes of interest (Ext1, NDST1, HS6ST2). We first produced knockout cell lines, transiently transfecting the gRNA/Cas9 plasmids (so that it would eventually be lost), and culturing them for 10 days. We then used FACS to identify and amplify cells that failed to take up fluorescently-tagged tau aggregates, assuming that this phenotype correlated with gene knockout. To rescue gene knockouts, we treated the sorted cells with lentivirus for 3 days to drive cDNA expression prior to uptake experiments (Figure 9). We completely reversed the knockout phenotype with cDNA overexpression (Figure 9). To further confirm the validity of the hits, we knocked out the genes of interest in the tau RD P301S biosensor cell line and exposed cells to tau aggregate seeds. Knockout of all 3 genes strongly reduced seeding activity (Figure 10).

**Figure 9:**
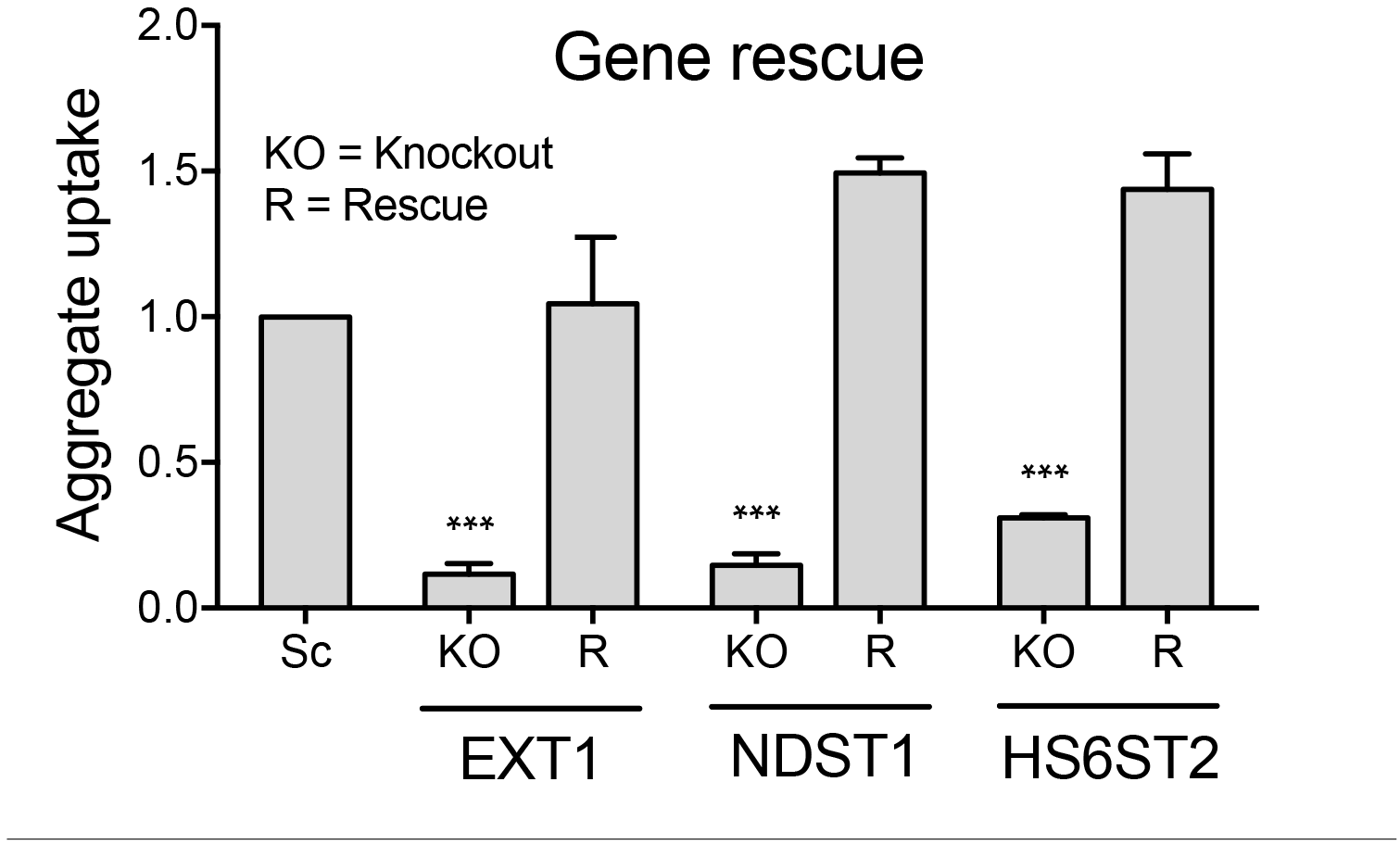
Rescue of tau uptake after gene knockout. We generated knockout cell lines for the main genes of interest (Ext1, NDST1 and HS6ST2) by transfection of the gRNA plasmids and FACS sorting. Knockout (KO) was completely reversible for all 3 genes when rescued (R) with cDNA lentivirus. Data was collected from two different experiments in triplicate for each cell line. The uptake in knockout cell lines was normalized to the uptake detected in scrambled control cells (Sc). Error bars show SD. For statistical analyses, we combined the averages for each experiment. ***P < 0.001; One-way Anova, Bonferroni.

**Figure 10:**
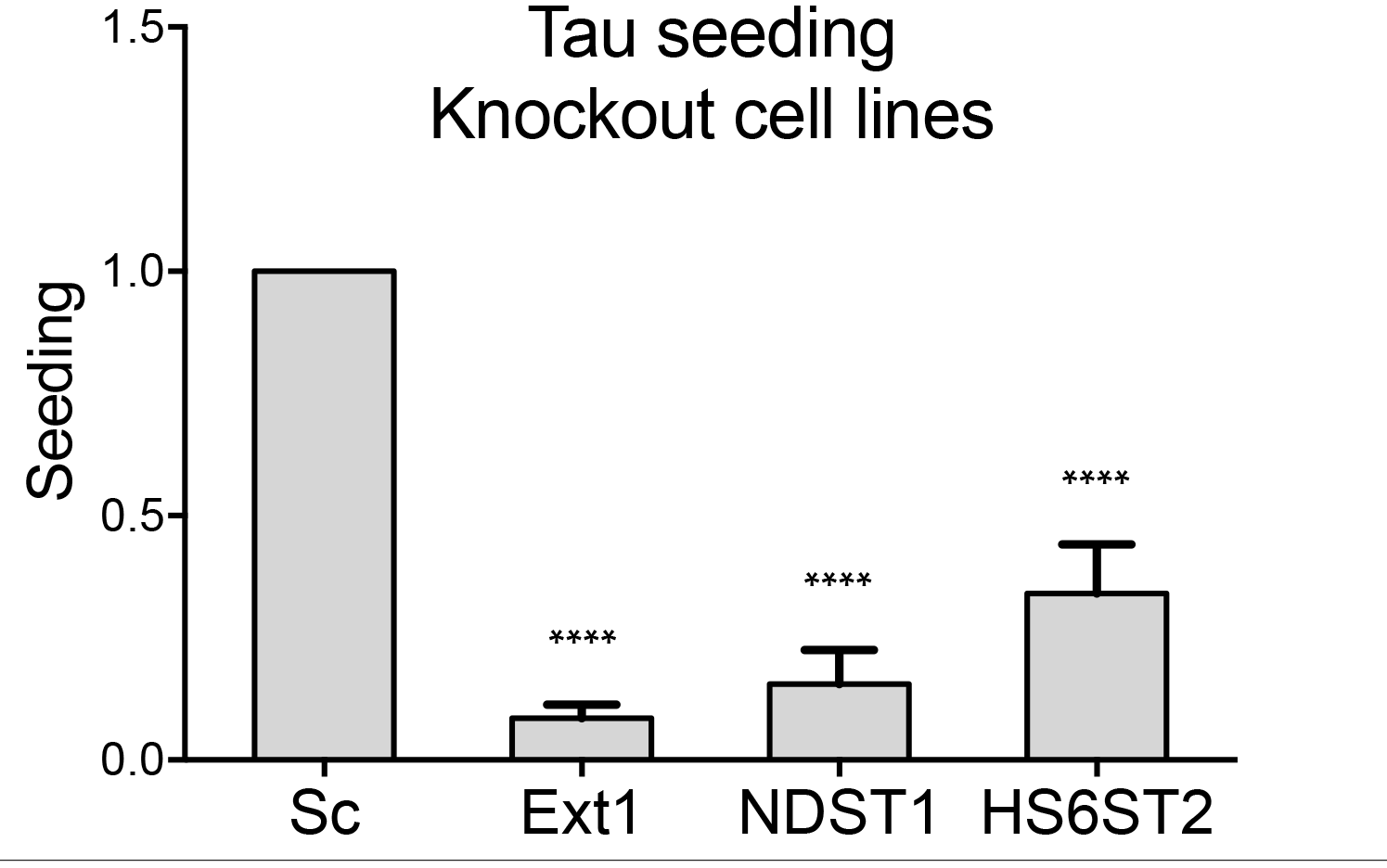
Inhibition of seeding by gene knockout. We generated knockout cell lines for the main genetic hits identified in our screen (EXT1, NDST1 and HS6ST2) and tested them for tau seeding by exposure to tau fibrils. We measured seeding by quantifying the percentage of FRET positive cells using flow cytometry. Knockout of the genes significantly reduced tau seeding at 48 hours. Data was collected in triplicate for each cell line and averaged over four experiments. The seeding in knockout cell lines was normalized to the seeding detected in scrambled control cells (Sc). Error bars show SD, using the average for each experiment. ****p < 0.0001; One-way Anova, Bonferroni.

In summary, targeted genetic knockdown of the HSPG biosynthesis pathway corroborates our previous data and implicates that tau uptake is specifically mediated by HSPGs and dependent on discrete sulfotransferases, while α-synuclein internalization seems to be more dependent on overall sulfation.

## Discussion

Using a combination of biochemical and cell-based assays we have investigated the structure-function relationships that govern the uptake of tau, α-synuclein and Aβ aggregates via HSPGs. The aggregate:GAG interaction, especially for tau, is determined by the pattern of sulfate moieties and the GAG length. We corroborated our findings with a genetic candidate screen of the HSPG synthesis pathway, and identified specific sulfotransferases that play a role in tau and α-synuclein uptake. Our data suggest that the interaction between HSPGs and aggregate seeds may be a regulated and specific biological process.

### Distinct sulfation requirements for aggregate binding

We originally employed a library of modified heparin polysaccharides spotted on a microarray to determine that the binding of tau aggregates depends on 6-O- and N-sulfation, whereas α-synuclein and Aβ aggregates require overall sulfation. We tested predictions from these studies in cell-based uptake and seeding assays, using heparins as competitive inhibitors of aggregate binding to the cell surface. N- and 6-O-desulfated heparins poorly inhibited tau uptake, whereas 2-O-desulfated heparin did not. This indicates a requirement for N- and 6-O sulfation for tau to bind heparin. We note that Zhao and colleagues recently described a requirement of 6-O sulfation for tau binding to GAGs in vitro (24), and Rauch and colleagues observed the relevance of 6-O-sulfation for cellular tau internalization in iPS-derived neurons and mouse brain slice culture (25). Differences in the observed importance of N-sulfation could be due to the different tau preparations used. On the contrary, removal of all 3 sulfate moieties reduced the inhibitory effect of the heparins for α-synuclein and Aβ aggregate uptake without regard to position. A recent report by Ihse and colleagues similarly found that the general degree of sulfation drives cellular α-synuclein uptake rather than specific modifications within the GAG chain (26). In summary, the required sulfation patterns for the GAG:seed interaction differ for distinct proteins: tau aggregates require specific sulfate moieties (N- and 6-O-sulfation), whereas α-synuclein and Aβ aggregate binding and uptake is mediated by overall sulfation and could be less specific. We finally note that sulfation requirements extend to viral uptake: baculovirus requires similar N- and 6-O-sulfation, but not 2-O-sulfation, for binding and entry into mammalian cells (27). This may imply that viruses enter cells by mechanisms that originally evolved to mediate aggregate transfer between cells.

### GAG chain length is critical

GAG chain length appears critical to mediate aggregate binding and uptake. Short heparin fragments (4-mer and 8-mer) exhibited no inhibitory activity for tau, while the longer fragments (12-mer and 16-mer) had only low inhibitory activity. Aβ seeds had low sensitivity for the small GAG sizes and an increasing sensitivity for longer fragments. α-Synuclein seeds displayed dose-dependent inhibition for all fractionated heparins. We conclude that chain length requirements for binding will vary among amyloid proteins.

### Specific sulfotransferases required for uptake

After screening all HSPG candidate genes, we observed that knockout of only five genes strongly reduced tau aggregate uptake. We previously validated EXT1 as a modifier of tau uptake (1). EXT2, EXTL3, NDST1 and HS6ST2 are unidentified genes in the context of tau pathology. EXT2 and EXTL3 have similar functions to EXT1, whereas NDST1 mediates N-sulfation, and HS6ST2 mediates 6-O-sulfation. Knockout of HS2ST1, the sole sulfotransferase of 2-O sulfation, did not alter tau seed internalization. On the contrary, knockout of EXT1, EXT2 and EXTL3, and NDST1, but not HS6ST2 reduced α-synuclein uptake. Knockout of HS2ST1 slightly reduced α-synuclein uptake. Altogether, the data corroborates our microarray and pharmacological data that tau uptake is mediated by on N- and 6-O-sulfation, whereas α-synuclein uptake depends on overall sulfation.

No other N- and 6-O-sulfotranferases were required for tau uptake. Some of the paralogs (e.g. NDST3, NDST4, and HS6ST3) have low or non-detectable expression levels in HEK293 cells (The Human Protein Atlas, (28)). It is possible that each paralog possesses unique substrate specificities and is not compensated by other members of the gene family (e.g. NDST2, HS6ST1). In addition, the expression levels of the paralogs might vary between cell and tissue types. Thus, in future studies, it will be crucial to investigate which enzyme paralogs drive N- and 6-O-sulfation in different cell and tissues types. We also recognize that future work must evaluate the requirements for these genes in the context of mammalian brain.

## Conclusion

Our results suggest that further studies to investigate the role of HSPGs and the relevance of structural requirements for aggregate uptake *in vivo* could be fruitful. A large body of literature indicates that sulfated GAGs can inhibit the infectivity of prion protein in vitro and in vivo (8,29–31), and our data suggest that this might apply to other amyloids as well. We are hopeful that the glycochemical principles described here might guide future drug development to block seed uptake and propagation of pathology in neurodegenerative disorders.

## Materials and Methods

### Protein preparation and labeling

Recombinant full-length wild-type tau and α-synuclein were purified and fibrillized as described previously (1). Huntingtin protein (Exon1 Q50) was prepared by solid-phase synthesis by the Keck Biotechnology Resource Laboratory at Yale University and was labeled and fibrillized as described previously (1). Synthetic Aβ42 was purchased by Anaspec unlabeled or with a biotin tag on the N-terminal amino acid. Biotin-Aβ42 was reconstituted in 1% NH_4_OH, lyophilized, and stored at −80°C. Unlabeled Aβ42 was reconstituted in 1% NH_4_OH, diluted in PBS, and stored at −80°C. Fibrillization of Aβ42 was achieved by bringing the protein concentration to 100 uM (Biotin Aβ42) or 110 uM (unlabeled Aβ42) in 10mM HCl and incubating for 24 hrs at 37°C. α-Synuclein and Aβ42 fibrils were dialyzed overnight into PBS. For uptake assays, tau, α-synuclein and Aβ42 were labeled with Alexa Fluor 647 succinimidyl ester dye (Invitrogen), quenched with 100 mM glycine and dialyzed overnight into PBS using dialysis cassettes (Thermo Scientific) with a molecular cutoff of 3500 Da (tau) or 2000 Da (α-synuclein and Aβ42) to remove excess dye molecules. The labeled fibrils were stored at 4°C.

### Heparin polysaccharide suppliers

The first experiment with microarrays was performed with heparins from Neoparin (Alameda, CA), which went out of business upon completion of the first part of our studies. Therefore, the heparins for the following studies were purchased from AMSbio (Cambridge, MA). The glycochemistry of these molecules is complicated and the quality of the products depends on the origin of the raw material used, the synthesis methods and the final purity of the preparations. We noticed differences in inhibitory activity when performing uptake assays with the same heparin from 2 different companies. However, we still derived qualitatively similar results, e.g. that certain defined moieties such as N-sulfation and 6-O sulfation are crucial for tau uptake, whereas overall charge is mostly driving the GAG:seed interaction for α-synuclein and Aβ. Also, not all the heparins used in the microarrays were available from AMSbio, which limited the number of heparins that could be tested in the second part of this study.

### Carbohydrate microarrays

As previously described (12), solutions of the polysaccharides (10 μl well in a 384-well plate) were spotted onto PLL-coated slides by using a Microgrid II arrayer (Biorobotics; Cambridge, UK) at room temperature and 50% humidity. Solution concentrations ranged from 0.5 μM to 15 μM, and the arrayer printed 1 nl of each concentration ten times. The resulting arrays were incubated in a 70% humidity chamber overnight and then stored in a low-humidity, dust-free desiccator. For protein binding studies, the slides were blocked with 3% bovine serum albumin (BSA) in PBS (5 ml) with gentle rocking at 37°C for 1 hr. Recombinant tau or α-synuclein or synthetic Aβ42 or huntingtin fibrils were reconstituted in 1% BSA in PBS, added to the slides in 50 μl quantities at a concentration of 1-2 μM, and incubated in a humidity chamber for 1 hr. The slides were then washed five times for 3 min each in PBS (5 ml) while gently rocking. After the washes, the slides were incubated with an anti-biotin IgG antibody conjugated to Cy5 in the dark with gentle rocking for 1 hr (1:5000, Molecular Probes), washed with PBS followed by H_2_O, and then dried under a gentle stream of N_2_. All incubations and washes were carried out at room temperature unless otherwise noted. The microarrays were analyzed at 635 nm for Cy5 by using a GenePix 5000a scanner, and fluorescence quantification was performed using GenePix 6.0 software with correction for local background. Each protein was analyzed in triplicate, and the data represent an average value for 10 spots at a given carbohydrate concentration. A similar experiment was performed with monomeric tau, α-synuclein, Aβ and huntingtin (not shown), but no binding could be detected (data not shown).

### Uptake assay for heparin experiments

C17.2 cells were plated at 4,000 cells per well in a 96-well plate. Fluorescently labeled aggregates (50 nM) were sonicated for 30 s at an amplitude of 65 (corresponding to ~80 Watt; QSonica) and pre-incubated overnight at 4°C in media containing the heparins at 4 different concentrations (0.2 - 200 μg/ml). The following morning, the aggregate-heparin complexes were applied to cells for 4 hours (tau and α-synuclein) or 20 hours (Aβ) for internalization in media vehicles of 150 ul per well. Incubation times were optimized empirically. Cells were harvested with 0.25% trypsin for 5 min and resuspended in flow cytometry buffer (HBSS plus 1% FBS and 1 mM EDTA) before flow cytometry. Cells were counted with the LSRFortessa SORP (BD biosciences). We determined the median fluorescence intensity (MFI) per cell to quantify cellular seed internalization. Each experiment was conducted 3 independent times with technical triplicates per condition, and a minimum of 2,000 single cells were analyzed per replicate. Conditions with single cell counts below 2,000 were excluded from analysis. We determined the average MFI of the replicates for each condition and standardized to seed uptake without inhibitor treatment within each experiment. The standardized averages of each condition were then combined for the graphs shown in this paper. Data analysis was performed using FlowJo v10 software (Treestar Inc.) and GraphPad Prism v7 for Mac OS X.

## Seeding assays

### Tau

Stable monoclonal FRET biosensor cell lines, expressing tau repeat domain (RD) with the aggregation-prone mutation P301S and tagged with either CFP or YFP, were previously generated in our laboratory (18,19). The seeding assay was conducted as described except that biosensor cells were plated at a density of 15,000 cells/well in a 96-well plate. Recombinant tau fibrils (100 nM) were sonicated for 30s at an amplitude of 65 (corresponds to ~80 Watt, QSonica) prior to use. Recombinant fibrils (100 nM) were pre-incubated overnight at 4°C in media containing the heparins at 4 different concentrations (0.2 – 200 μg/ml). At 20-30% confluency, the seed-heparin complexes were applied to the cells in media volumes of 50 ul per well, and cells were incubated for an additional 48 hrs. Contrary to previous publications, we did not use lipofectamine to drive internalization of seeds since we intended to monitor HSPG-mediated uptake into the cells and the inhibitory activity of heparin blocking this process. The cells were harvested with 0.05% trypsin and post-fixed in 2% paraformaldehyde for 10 min, then resuspended in flow cytometry buffer (HBSS plus 1% FBS and 1 mM EDTA). The LSRFortessa SORP (BD Biosciences) was used to perform FRET flow cytometry. We quantified FRET as previously described (18,19) with the following modification: We identified single cells that were YFP-and CFP-positive and subsequently quantified FRET positive cells within this population. For each data set, 3 independent experiments with 3 technical replicates were performed. For each experiment, a minimum of approximately 10,000 single cells per replicate was analyzed. Wells with YFP-/CFP-positive cell counts below 5,000 cells were excluded from the analysis. Data analysis was performed using FlowJo v10 software (Treestar Inc.) and GraphPad Prism v7 for Mac OS X.

### α-Synuclein

A stable monoclonal FRET biosensor cell line was previously established in our laboratory, expressing full length α-synuclein with the disease-associated mutation A53T, and tagged either with CFP or YFP (18,19). The seeding assay was performed similar to the tau seeding assay, except that biosensor cells were plated at a density of 10,000 cells/well in a 96-well plate. Recombinant α-synuclein fibrils (200 nM) were sonicated for 30s at an amplitude of 65 (corresponds to ~80 Watt, QSonica) prior to use. The fibrils were pre-incubated overnight at 4°C in media containing the heparins at 4 different concentrations (0.2 – 200 μg/ml). At 20% confluency, the seed-heparin complexes were applied to the cells in media vehicles of 50 ul per well, and cells were incubated for 48 hrs. Cells were then split 1:5 into a new 96-well plate and incubated overnight. The next day, a second treatment with α-synuclein fibril-heparin complexes was performed similar to above. After an additional 48 hrs, cells were harvested for flow cytometry. For the data shown in this paper, we performed 2 independent experiments, with 3 technical replicates of each condition in each case. Data analysis was performed as described above for tau seeding.

### Candidate screen using CRISPR/Cas9 knockout and lentiviral transduction

5-6 human gRNA sequences per gene were derived from the GeCKO v2 libraries (20). For all gRNA sequences not commencing with guanine, a single guanine nucleotide was added at the 5′ end of the sequence to enhance U6 promoter activity. DNA oligonucleotides were synthesized (IDT DNA) and cloned into the lentiCRISPRv2 vector (20) for lentivirus production. The plasmids for 5-6 gRNAS for each gene were pooled together. Lentivirus was created as described previously (32): HEK293T cells were plated at a concentration of 1×10^6^ cells/well in a 6-well plate. 18 hours later, cells were transiently co-transfected with PSP helper plasmid (1200 ng), VSV-G (400 ng), and gRNA plasmids (400 ng) using 7.5 μL TransIT-293 (Mirus). 72 hours later, conditioned media was harvested and centrifuged at 1000 rpm for 5 minutes to remove dead cells and debris. Lentivirus was concentrated 50x using lenti-X concentrator (Clontech) with the concentrated pellet being re-suspended in PBS with 25 mM HEPES, pH 7.4. For transduction, a 1:25 to 1:40 dilution of virus suspension was added to HEK293T cells or Tau RD P301S biosensor cells at a cell confluency of 20% in a 96 well plate. 20 hrs post-transduction, infected cells were treated with 2 μg/mL of puromycin (Life Technologies) and cultured for additional 2 days, followed by passaging 1:5 and a second round of virus and puromycin application. The cells were kept in culture for at least 10 days after the first lentiviral transduction before using them for uptake and seeding experiments. After identification of the genes of interest, we made knockout cell lines using the available gRNAs individually to identify the functional gRNAs. We identified 2-3 gRNAs per gene and used these for further knockout experiments. In addition, we used a single functional gRNA for each gene to produce knockout cell lines for analysis by Tracking of Indels by Decomposition (TIDE)(21) to confirm the presence of indels in the predicted DNA regions.

### Uptake assay for knockout screen

The uptake assay for the knockout screen was conducted as described above with the following modifications: Knockout cell lines were plated at 20,000 cells per well in a 96-well plate. The next day, aggregates labeled with Alexa Fluor 647 were sonicated for 30 s at an amplitude of 65 (corresponds to ~80 Watt, QSonica), mixed to transferrin labeled with Alexa Fluor 488 (Life Technologies) and applied to the cells in a vehicle of 50 ul media for 4 hours. Final concentrations of tau and α-synuclein in the media were 25 nM; of transferrin 25 μg/ml. Cells were harvested with 0.25% trypsin for 5 min and re-suspended in flow cytometry buffer (HBSS plus 1% FBS and 1 mM EDTA) before flow cytometry. Cells were counted with the LSRFortessa SORP (BD biosciences). Data was collected from the same cell lines on two different days in duplicates. For each replicate, approximately 10,000 single cells were analyzed. Uptake was normalized to the uptake detected in scrambled control cells (Sc). For the analysis, we combined the averages for each experimental day to determine the multiplicity adjusted P values using one-way Anova and Dunnett testing. Data analysis was performed using FlowJo v10 software (Treestar Inc.) and GraphPad Prism v7 for Mac OS X.

### Rescue experiments

2-3 functional gRNA plasmids identified as described above were pooled and used for transfection. Briefly, 60,000 HEK293T cells were plated per well in a 24 well plate. The next day, 500 ng DNA was combined with OptiMEM (Gibco, Life Technologies) to a final volume of 25 ul. Similarly, 2.5 ul of lipofectamine-2000 (Invitrogen, Life Technologies) was combined with OptiMEM (Gibco, Life Technologies) to a final volume of 25 ul. Both solutions were mixed after 5 min of incubation and incubated for additional 20 min at room temperature, followed by drop-wise addition to the cells at about 50% of cell confluency. We applied puromycin once at 2 ug/ml 24 hours after transfection. Cells were maintained at least 10 days prior to an uptake experiment with Alexa Fluor 647 labeled tau fibrils and FACS sorting for Alexa Fluor 647 negative cells. The FACS sorted populations were re-plated in a 96 well plate and maintained until further use. Ext1 cDNA was subcloned from pCMV3-Ext1-HA (Sino Biological) into lentiviral FM5 vector (33). NDST1 cDNA was subcloned from pUC75-NDST1 (Genscript) into FM5:HA vector (33). HS6ST2 cDNA was subcloned from pANT7-cGST-HS6ST2 (DNASU Plasmid depository, Arizona Stat University (34)) into lentiviral FM5:HA vector (33). All plasmids were sequenced before use. Lentivirus was produced and concentrated 100x as described above. Lentivirus was applied at a 1:150 to 1:75 dilution to the knockout cell lines for three days prior to uptake experiments. Data was collected from the same cell lines on two different days in triplicates. For each replicate, approximately 5,000 single cells were analyzed. Uptake was normalized to the uptake detected in scrambled control cells (Sc). For the analysis, we combined the averages for each experimental day to determine the multiplicity adjusted P values using one-way Anova and Bonferroni testing. Data analysis was performed using FlowJo v10 software (Treestar Inc.) and GraphPad Prism v7 for Mac OS X.

### Seeding assay in knockout cell lines

We generated knockout cell lines as described above for the genes of interest in our P301S FRET biosensor line. For seeding experiments, 100 nM of recombinant tau fibrils were applied for 48 hours prior to flow cytometry. Data was collected from the same cell lines on four different days in triplicates. For each replicate, approximately 10,000 single cells were analyzed. Seeding was normalized to the seeding detected in scrambled control cells (Sc). For the analysis, we combined the averages for each experimental day to determine the multiplicity adjusted p values using one-way Anova and Bonferroni testing. Data analysis was performed using FlowJo v10 software (Treestar Inc.) and GraphPad Prism v7 for Mac OS X.

## Acknowledgements

This work was supported by the Tau Consortium (M.I.D); NIH grant 1F31NS079039 (B.B.H.); grant “Rotation Program for Junior Researchers” by the Faculty of Medicine, RWTH Aachen University (B.E.S.).

